# Analysis of selection methods to develop novel phage therapy cocktails against antimicrobial resistant clinical isolates of bacteria

**DOI:** 10.1101/2020.10.13.337345

**Authors:** Melissa EK Haines, Francesca E Hodges, Janet Y Nale, Jennifer Mahony, Douwe van Sinderen, Joanna Kaczorowska, Bandar Alrashid, Mahmuda Akter, Nathan Brown, Dominic Sauvageau, Thomas Sicheritz-Ponten, Anisha M Thanki, Andrew D Millard, Edouard E Galyov, Martha RJ Clokie

**Author notes:** Corresponding Author – Melissa Haines.

## Abstract

Antimicrobial resistance (AMR) is a major problem globally. The main bacterial organisms associated with urinary tract infection (UTI) associated sepsis are *E. coli* and *Klebsiella* along with *Enterobacter* species. These all have AMR strains known as ESBL (Extended Spectrum Beta-Lactamase), which are featured on the WHO priority pathogens list as ‘critical’ for research. Bacteriophages (phages) as viruses that can infect and kill bacteria, could provide an effective tool to tackle these AMR strains.

There is currently no ‘gold standard’ for developing a phage cocktail. Here we describe a novel approach to develop an effective phage cocktail against a set of ESBL-producing *E. coli* and *Klebsiella* largely isolated from patients in UK hospitals. By comparing different measures of phage efficacy, we show which are the most robust, and suggest an efficient screening cascade that could be used to develop phage cocktails to target other AMR bacterial species.

A target panel of 38 ESBL-producing clinical strains isolated from urine samples was collated and used to test phage efficacy. After an initial screening of 68 phages, six were identified and tested against these 38 strains to determine their clinical coverage and killing efficiency. To achieve this, we assessed four different methods to assess phage virulence across these bacterial isolates. These were the Direct Spot Test (DST), the Efficiency of Plating (EOP) assay, the planktonic killing assay and the biofilm assay.

The final ESBL cocktail of six phages could effectively kill 23/38 strains (61%) for *Klebsiella* 13/19 (68%) and for *E. coli* 10/19 (53%) based on the planktonic killing assay data. The ESBL *E. coli* collection had six isolates from the prevalent UTI-associated ST131 sequence type, five of which were targeted effectively by the final cocktail. Of the four methods used to assess phage virulence, the data suggests that planktonic killing assays are as effective as the much more time-consuming EOPs and data for the two assays correlates well. This suggests that planktonic killing is a good proxy to determine which phages should be used in a cocktail. This assay when combined with the virulence index also allows ‘phage synergy’ to inform cocktail design.

## Introduction

Antimicrobial resistance (AMR) is a major global challenge. It is part of the key target priorities for several prominent organisations including the World Health Organisation (WHO), European Centre for Disease Prevention and Control (ECDC) and National Institute of Health Research (NIHR) (Tacconelli et al., 2018). It has been predicted that more people will die of AMR than cancer by 2050 and AMR associated deaths are estimated to be approximately 10 million people per year (O’Neill, 2014). AMR has been compounded by a reduction in novel antibiotic discovery, the persistent use of antibiotics and thus, the rapid emergence of bacterial strains that are resistant to⍰both existing and new antibiotics (Tacconelli et al., 2018). The most clinically relevant group of multi-drug resistant (MDR) pathogens are referred to collectively as the ESPAKEE organisms (Gram-positive⍰<u>E</u>nterococcus faecium⍰and⍰<u>S</u>taphylococcus aureus, as well as Gram-negative⍰<u>P</u>seudomonas aeruginosa, <u>A</u>cinetobacter baumannii, *<u>K</u>lebsiella* pneumoniae, Enterobacter⍰species and Escherichia coli), and are together responsible for the majority of hospital-acquired infections (Pendleton et al., 2013). Urinary tract infections (UTIs) are prevalent and can cause serious infections *per se* but can also act as infection sources for sepsis (urosepsis) and septicaemia. The majority of organisms associated with urosepsis are *E. coli*, which are responsible for 50% of cases, and *Klebsiella* along with other Enterobacter species, which total 15% of cases (Kalra and Raizada, 2009). Furthermore biofilm formation has been shown to be crucial in infections such as catheter-associated UTIs with both *E. coli* and *Klebsiella* (Hancock et al., 2010).

Extended Spectrum Beta Lactamases (ESBL) are plasmid-mediated enzymes that, if expressed by a bacterial strain, confer resistance to antibiotics containing a beta-lactam ring in their molecular structure such as penicillins, cephalosporins and carbapenems (Livermore, 1987; Sykes and Matthew, 1976; van Duin and Doi, 2017). ESBL-producing strains⍰of both E.⍰coli⍰and⍰*Klebsiella* have been detected from a variety of sources worldwide (Bush, 2018).⍰They pose a serious global public health threat due to the difficulties associated with treatment of infections with ESBL-producing bacteria. Although, Sakellariou et al. (2016) reported no difference⍰in mortality⍰rates of infections⍰caused by either⍰ESBL-producing⍰*E. coli*⍰(23.8%)⍰or⍰ESBL-producing⍰*Klebsiella*⍰(27.1%), they did report that septicaemia associated with⍰ESBL-producing⍰*Klebsiella*⍰has a higher morbidity (sepsis with organ failure).⍰

The⍰decline in⍰antibiotic⍰discovery and emergence⍰of resistance⍰to last line antibiotics⍰(Pendleton et al., 2013),⍰motivates the need for⍰alternative antimicrobials. A promising solution is the therapeutic application of bacteriophages (phages), which are viruses that kill bacteria. Phage therapy has a long history of use in countries such as Georgia,⍰Poland and France⍰(Ansaldi et al., 2018; Gorski et al., 2018; Kutateladze and Adamia, 2008)⍰where it has been used⍰alongside or instead of antibiotics to treat bacterial infections for⍰more than⍰80 years.⍰⍰There is a critical need to widen access to this therapy,⍰either as an alternative to or as a supplement for antibiotic treatment. If phage therapy is to be developed in the Western world, it is advisable to focus on bacterial diseases for which no other treatments exist and those which have high levels of AMR⍰(Tomas et al., 2018). A phage cocktail is a mixture of several⍰phages and has two potential clinical advantages (2012). One is to combine the individual⍰phages⍰to broaden the number of strains the phages are able to infect. The second is to combat resistance, which can occur with the use of single⍰phages.⍰By using a cocktail of phages, strains that become resistant to one phage can be targeted by other⍰phages⍰within the cocktail.⍰In the context of the current study, the primary goal for the phage cocktail was to provide a broader host range than any of the individual⍰phages⍰alone. Host range coverage was prioritised over efficiency of killing with regards to the phage cocktail selection. This is because in a clinical context, it would be beneficial to provide partial treatment to a wider number of patients, allowing synergy with the immune system and antibiotics, rather than treating only a select few patients⍰(Chan et al., 2013; Mattila et al., 2015). The overall aim was to identify⍰phages⍰that individually have broad host ranges and collectively when combined would cover ~90% of the either the ESBL-producing⍰*E. coli*⍰or⍰*Klebsiella*⍰collection.

Although phage cocktails have been designed and their efficacy reported in the literature previously, there are no current guidelines to standardise the development of an optimised cocktail for resistant bacteria or indeed to predict the efficacy of⍰phages⍰at least under⍰*in vitro*⍰conditions. Through the development of the phage cocktail in the current study, we have generated a data set that allows comparison of four different methods of assessing phage virulence across a panel of 38 bacterial isolates. These tests⍰are:⍰ Direct Spot Test (DST), Efficiency of Plating (EOP) assay, a planktonic killing assay and a biofilm assay. Both the DST and EOP assay are frequently utilised in the determination of phage virulence (Mirzaei and Nilsson, 2015) and both tests use the double agar plate method. The DST is a reasonably good method for initial host range screening, but it does not provide a reliable indication that the phage can replicate on the host strain. The EOP assay indicates productive infection of the host strain by the phage from which the efficiency of infection of the host can be determined. The planktonic killing assay was assessed as an alternative to the labour-intensive DST and EOP approaches. This method monitors the optical density⍰of a liquid culture of bacteria⍰to which a⍰phage combination was added⍰using a plate reader over 24 hours. The previous three methods examine the virulence of⍰phages⍰based on killing bacteria under ‘normal’ growth conditions⍰in vitro and so the final method chosen was a biofilm assay. This assay provides an insight into phage virulence in an⍰in-vitro⍰model of infection and biofilm formation. Genomic analysis was performed on all 38 ESBL-producing⍰clinical isolates to determine the relationships between susceptibility to phage infection and genomic content. The genetic relationship between the most sensitive and most resistant clinical isolates was determined.⍰ The final six⍰phages⍰selected for the ESBL phage cocktail were also sequenced to confirm suitability for phage therapy and ensure they did not encode for any known undesirable traits.⍰

This article focuses on the development of a phage cocktail that is effective against ESBL-producing⍰*E. coli*⍰and⍰*Klebsiella*⍰that were largely isolated from patients in UK hospitals. In producing this data, we describe an efficient screening cascade to develop cocktails, which will be relevant for other target AMR bacterial species.⍰ This data⍰shows⍰a novel, direct⍰comparison⍰of results across⍰the four phage virulence tests for⍰individual clinical isolates and indicates that⍰the⍰planktonic killing assays are a reliable and time efficient way to⍰assess phage⍰efficacy.⍰

## Materials & Methods

### Bacterial strains

38 strains of ESBL-producing bacteria were examined during this study; 19 *E. coli* and 19 *Klebsiella*. All strains were clinical isolates from urinary tract infections; 14 of the *E. coli* isolates were from Leicester Royal Infirmary, UK; 5 from Huashan Hospital, Shanghai and 19 *Klebsiella* isolates from Leicester Royal Infirmary, UK [Supplementary Data Table S1]. All bacteria were grown at 37°C in either Luria-Bertani Broth (LB - Thermo Fisher Scientific, United Kingdom) at 100 rpm or on LB 1% (w/v) agar plates. All strains were stored in 50% glycerol stocks at −80°C until required. The bacterial strains were sequenced by MicrobesNG with the Standard Whole Genome Service, Illumina Sequencing by sending the bacterial strains as samples.

### Phage Collection, Isolation, Amplification and Visualisation

Phages were collated from several sources with the majority coming from collaborations with other research projects [Supplementary Data Table S2]. Phages were isolated using the method previously described by (Kropinski et al., 2009). To identify phages, 100 μl enrichment, 100 μl culture and 3 ml LB 0.5% (w/v) agar were poured onto a LB 1% (w/v) agar plate and incubated overnight at 37°C. Single plaques were picked and transferred to 500 μl SM Buffer (100mM NaCl (Sigma-Aldrich, United Kingdom), 8mm MgSO_4_•7H2O (Sigma-Aldrich, United Kingdom), 0.1% (w/v) gelatin (Sigma-Aldrich, United Kingdom), 50mM Tris-HCl pH 7.5 (Sigma-Aldrich, United Kingdom)). This process was repeated to give five rounds of single plaque purification and stored in SM buffer.

Phage stocks were made using the double layer agar method. Briefly, an overnight culture of the host strain was diluted 1:100 in LB and grown for 2 hours at an ~OD_550_ of 0.2 at 37°C, 100 rpm. 500 μl of the bacterial culture and 200 μl of phage stock were added to 8 ml of 0.5% (w/v) LB agar and poured onto 120×120 mm square LB 1% (w/v) agar plates. The plates were incubated overnight at 37°C. The plates were agitated for 2 hours in 10 ml SM buffer. The top layer was removed and centrifuged at 4,000 × g for 15 mins. The supernatant was filter-sterilised through 0.2 μm pore size filters and the resultant phage stock titre was determined using double agar overlay plaque assays (Kropinski et al., 2009). Stock was stored at 4°C. Phage UP17 (vB_EcoM_UP17) was propagated using *E. coli* EA2; phage JK08 (vB_SsoM_JK08) – *E. coli* MH10; phage 113 (vB_SsoM_113) – *Shigella sonnei* B31; phage 2811 (vB_KpnS_2811) – *Klebsiella pneumoniae* KR2811; phage 311F (vB_KpnM_311F) – *K. pneumoniae* KR311; phage 05F (vB_KpnM_05F) – *K. pneumoniae* MH05.

Transmission Electron Microscopy imaging for the phages UP17, 113, 2811, 311F and 05F was performed at University of Leicester, UK. The phages were negatively stained with 1% (w/v) uranyl acetate on 3 mm carbon-coated copper grids. Visualised with a JEM-1400 transmission electron microscope (JEOL UK Ltd., United Kingdom) with an accelerating voltage of 120 kV. Digital images were collected with an Xarosa digital camera (EMSIS, Germany) with Radius software for phage 113; all other phages were imaged using a Megaview III digital camera (EMSIS, Germany) instead. Imaging for phage JK08 was performed at the Max Rubner-Institut, Germany with with a Tecnai 10 transmission electron microscope (FEI, Eindhoven, The Netherlands) operated at an acceleration voltage of 80 kV.

### Direct Spot Testing (DST)

Bacterial cultures were grown overnight, then diluted 1/100 in LB and grown for 2 hours to ~OD_550_ of 0.2. 500 μL of the culture was added to 8 ml 0.5% (w/v) LB agar kept molten at 55°C and poured onto LB 1% (w/v) agar square 120×120 mm plates. 20 μl of phage stock (10^9^/10^10^) was spotted onto the plate, left to dry and then incubated overnight at 37°C. The appearance of the spot was graded: +++ complete lysis; ++ lysis with resistant colonies; + hazy lysis; 0 visual plaques [Supplementary Data S3 & S4].

### Efficiency of Plating (EOP)

This method has been previously described by Kutter, 2009; 5 mM calcium chloride was supplemented to the 0.5% (w/v) LB agar for the *E. coli* and *Shigella* phages. Plaques on each plate were counted and the relative EOP was given as the ratio between the phage titre in pfu/ml (plaque forming units/ml) for the test host strain and the titre of the propagating host strain. Propagating host for phages UP17, JK08 and 113 was *E. coli* MH10. The propagating host for 2811 – K. pneumoniae KR2811; for 311F – K. pneumoniae KR311; and for 05F – *K. pneumoniae* MH05.

### Planktonic Killing Assay

Experiments were carried out using the BMG Labtech SPECTROstar Omega, using a flat bottom 96 well plate (Sarstedt, Germany). 100 μl of a 1:100 dilution of overnight cultures was added to the 96 well plate, grown to A_600_ OD 0.15 (1×10^8^ CFU/ml), then 100 μl of phage cocktail (containing 1×10^8^ PFU/ml of each individual phage) was added. Working with a MOI of 1:1; throughout all the experiments. Final concentrations were achieved using LB as a diluent. The microtiter plates were securely sealed using parafilm M (Amcor, US). OD readings (A_600_) were taken every 5 mins for a total of 24 hours with shaking 10 seconds prior to each reading. The microtiter plate had a positive control for every individual clinical strain for comparison, as well as a negative control (LB only) and 3 blanks (LB and gentamicin 10 μg/ml). Each cocktail was repeated in triplicate for each ESBL-producing clinical isolate and the data was merged to give a single killing assay curve.

The killing assay curves were analysed by an objective method (Storms et al., 2019) which was devised using the generated curve to give a ‘virulence index.’ The virulence index score was calculated comparing the area under the curve of the individual phage or cocktail against the positive control whilst in log phase. This virulence index was normalised to a figure between 0 - 1, 0 = not effective and 1 = highly effective.

### Biofilm Assay

Bacterial cultures were grown overnight at 37°C, 100 rpm. 100 μl of 1:100 dilution in LB of each bacterial strain was added to 96 well flat bottom microtiter plate in triplicate for both controls and phage cocktail treated. The whole experiment was also repeated in triplicate for all bacterial strains. After 24 hours at 37°C, the LB was removed, and each well was washed with PBS. For control, 100 μl of fresh LB was added instead of 100 μl of final ESBL cocktail was added (10^8^ PFU/ml of each individual phage). After an additional 24 hours of incubation, 20μl of resazurin (0.15 mg/ml - Sigma-Aldrich) was added and incubated at 37°C. OD readings were taken at A_595_ with Labtech.com LT-4500 at 4 h and 24 h post incubation.

### Phage DNA extraction

Phage lysate at titres of 10^11^ PFU/ml was used to extract DNA using a modified phenol-chloroform-isoamyl method as previously described (Nale et al., 2015). The final DNA pellet was dissolved in 5 mM Tris HCl. This method only applies to phages UP17, 113, 2811, 311F and 05F. For phage JK08, DNA isolation was performed using the Norgen Phage DNA isolation Kit (Norgen Biotek, ON, Canada) according to the manufacturer’s instructions.

### Sequencing and Bioinformatic Analysis

Genome sequencing was conducted by MicrobesNG (http://www.microbesng.uk), which was supported by the BBSRC (grant number BB/L024209/1) for phages UP17, 311F and 05F as well as all the bacterial genomes. *De novo* assembly of the trimmed reads using Trimmomatic 0.30 (Bolger et al., 2014) from MicrobesNG was carried out using SPAdes genome assembler 3.12.0 (Bankevich et al., 2012) with default settings.

For the bacterial genomes, contigs were annotated using Prokka v1.12 (Seemann, 2014) and the assembly metrics were calculated using QUAST 5.0.2 (Gurevich et al., 2013). MLST 2.16.2 was used for characterisation of the bacterial strains (Seemann - https://github.com/tseemann/mlst). ABRicate with Resfinder database was used with default settings to screen the genome of each strain for the presence of antimicrobial resistance and virulence genes (Feldgarden et al., 2019; Zankari et al., 2012).

Sequence data for the bacterial genomes was also used to create phylogenetic trees (Figure 4) using MEGA7 v7180411 (Kumar et al., 2018) and visualised using iTOL v5.5 (Letunic and Bork, 2007) based on the core genome SNPs. For phages 113 and 2811, the genomes were sequenced using an Illumina MiSeq, with a v3 kit (600 cycles). Genomic libraries were prepared using the Illumina Truseq Nano DNA library Preparation Kit as per the manufacturer’s instructions. The genomes were assembled using MEGAHIT (Li et al., 2015); phage 2811 (version 1.2.1) and phage 113 (version 1.1.4). Phage termini were identified using PhageTerm v1.0.11 (Garneau et al., 2017). Phage JK08 was sequenced using an Illumina MiSeq using a v2 kit (2 × 250). Illumina Truseq PCR-free library preparation kit was used as per manufacturer’s instructions for genomic library preparation. Genome assembly was performed with MIRA v4.0.2 (Chevreux et al., 1999).

The genomes of phage UP17, 311F and 05F were assembled by subsampling reads to an approximate coverage of 100× with seqtk (https://github.com/lh3/seqtk) and assembled with SPAdes v3.12.0 with only-assembler option (Bankevich et al., 2012). Phage genomes were annotated as previously described (Michniewski et al., 2019). To check for antibiotic resistance and virulence genes within the phage genomes, ABRicate was used with the card and vfdb databases respectively.

### Accession Numbers

All bacterial and phage genomes were submitted to the European Nucleotide Archive (ENA) under project accession number PRJEB34549. Individual accession numbers are provided in Supplementary Data Tables S1 and S2.

### Statistical Analysis

GraphPad Prism 7.04 (La Jolla, CA, USA) was used for statistical analysis for the biofilm assays. The results were expressed as mean +/−SEM after analysis with 2-way ANOVA. A p-value < 0.05 was considered significant.

## Results

### Comparison of phage virulence methods – DST, EOP and planktonic killing assay

The three methods used to assess phage virulence: DST, EOP and the planktonic killing assay were compared. These three tests form the basis for the initial screening of a phage library to identify phages with the broadest host range. The data generated also allowed direct comparison of DST versus EOP, as these two assays are commonly used to characterise phages (Montso et al., 2019; Rivera et al., 2019; Sybesma et al., 2016) [Supplementary Data Table S5 & S6].

The final three phages selected for the final ESBL cocktail based on their effectiveness against the ESBL-producing *E. coli* strains were UP17, JK08 & 113. With phages 2811, 311F and 05F selected to target ESBL-producing *Klebsiella*. Phages were selected based on the results of the DST, EOP and planktonic killing assay data (Figure 1). The selection of the final three phages was based on combining the minimal number of phages to have the maximal effect. For example, with the *E. coli* phages using four phages resulted in the same percentage coverage of using only three [Supplementary Data Table S5].

**Figure 1.**
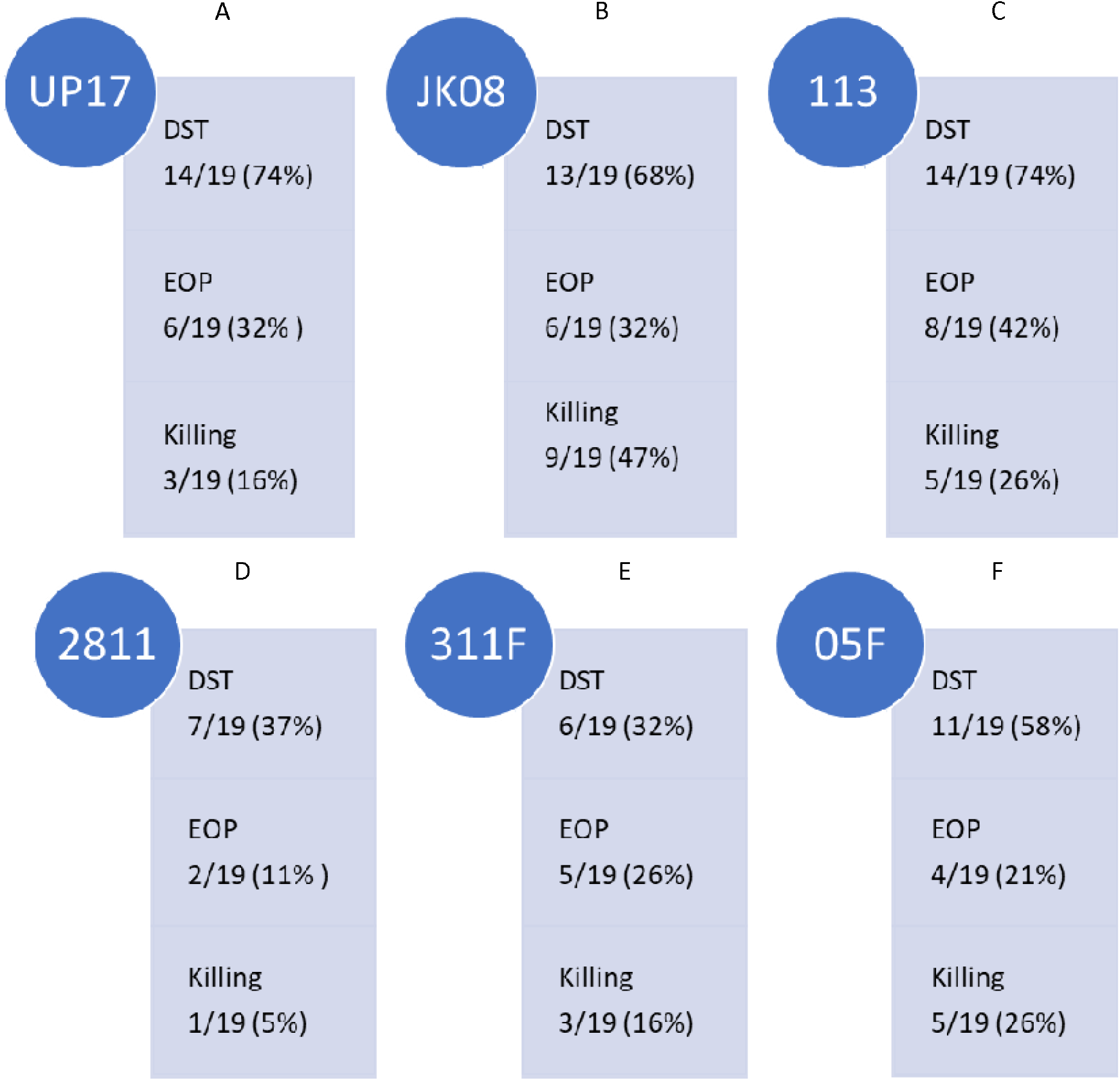
Summary of the ESBL-producing E.coli clinical isolate (n=19) coverage of final E.coli phages (A) UP17, (B) JK08 and (C) 113 and ESBL-producing *Klebsiella* clinical isolate (n=19) coverage of final *Klebsiella* phages (D) 2811, (E) 311F and (F) 05F across the three selection test (Direct Spot Test [DST], Efficiency of Plating [EOP] and Killing [Planktonic Killing Assay]). Isolate coverage was determined by the following parameters: DST ≥ + appearance score; EOP > 0.01; Killing ≥0.2 virulence index score.

The final three *E. coli* phages were selected on the basis they have the broadest clinical isolate coverage. The following coverage was observed: phages UP17, JK08 and 113 could lyse 14/19 (74%), 13/19 (68%) and 14/19 (74%) of *E. coli* clinical isolates, respectively (Figure 1). When the phages were combined based on DST data, they provided coverage of 18/19 clinical isolates (95%) [Supplementary Data Table S5]. The final three phages selected to be effective against the ESBL-producing *Klebsiella* clinical isolates gave overall coverage of 17/19 (89%) based on DST data [Supplementary Data Table S6]. In comparison the individual phages gave the following coverage: phage 2811 lysed 7/19 (37%), phage 311F lysed 6/19 (32%) and phage 05F lysed 11/19 (58%) (Figure 2).

**Figure 2.**
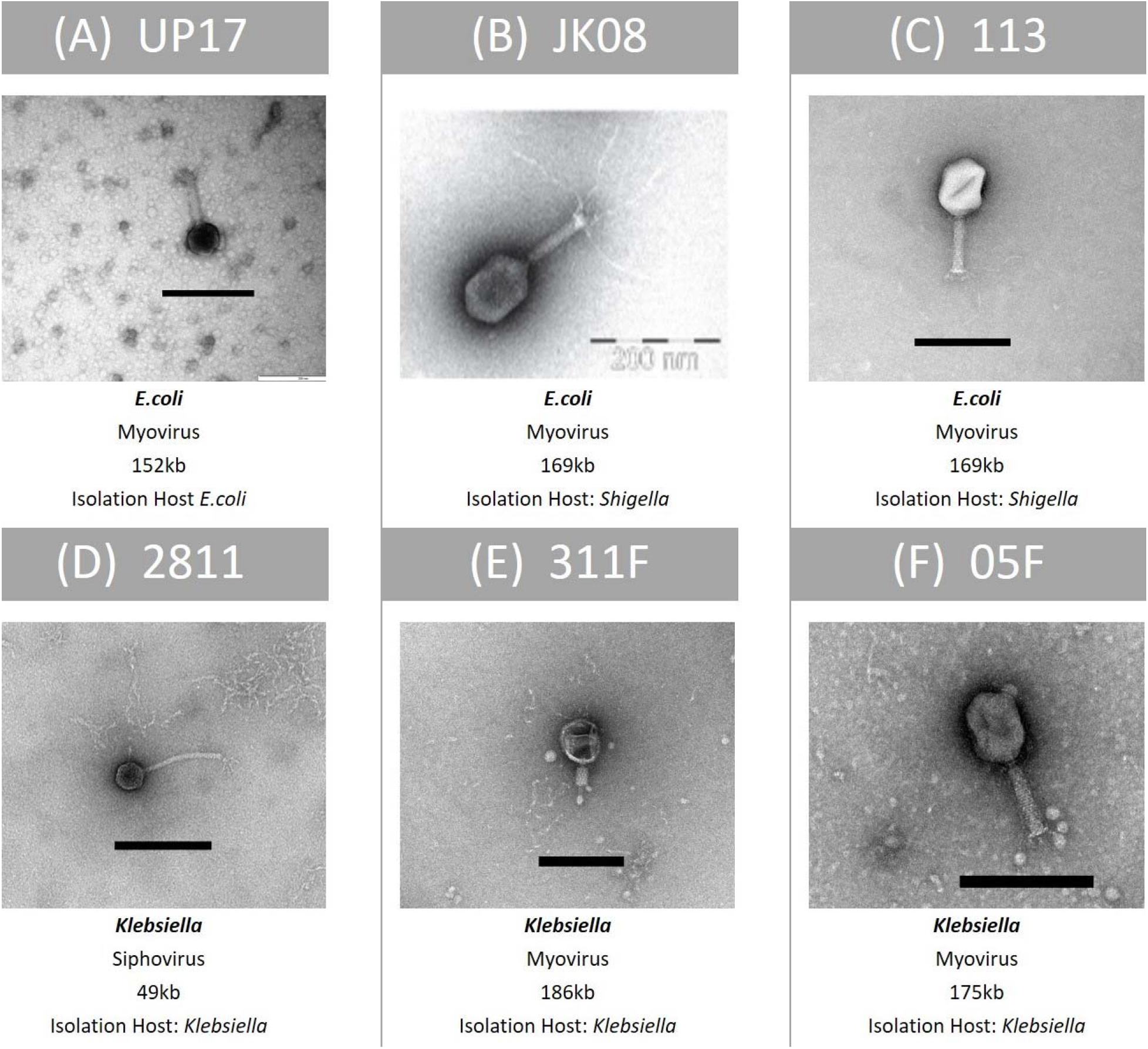
Summary of key features of the final six phages within the ESBL cocktail – TEM image, family classification, genome size and species of propagation host. From top row left to right, Panel (A): Phage UP17, Panel (B): Phage JK08, Panel (C): Phage 113. From bottom left to right: Panel (D): Phage 2811, Panel (E): 311F and Panel (F): Phage 05F. Black bar represents 200nm

The DST data highlighted which clinical isolates were lysed by the phages. To determine if the phages could efficiently replicate on the clinical isolates they infected, EOP studies were conducted. A detectable EOP was defined as the ratio compared to the control stain was > 0.01. Across all the phages, the number of isolates on which they could replicate within (EOP) were lower than those lysed (DST) (Figure 1). Collectively for the three *E. coli* infecting phages, EOP data showed 13/19 strains (68%) compared to 18/19 (95%) predicted by the DST. DST overestimates the efficiency of killing compared to EOP and planktonic killing assay. For example, UP17 only effectively replicates in 6/14 of the clinical isolates identified by DST.

There is a closer relationship between the planktonic killing assay and EOP data; but the trend appears to be that planktonic killing assay is lower than EOP isolate coverage. For example, the planktonic killing assay showed that phage 05F was effective (virulence index ≥ 0.2) for 5/19 (26%) clinical isolates compared with EOP 4/19 (21%) (Figure 1). Based on EOP data for phage 2811, it suggests that the phage could only replicate on 2/19 (11%) clinical isolates compared with 1/19 (5%) on the planktonic killing assay (Figure 1).

### Characterisation of the final six phages selected for the ESBL phage cocktail

The final phages selected to target ESBL-producing *E. coli* were UP17, JK08, 113 and for *Klebsiella* the final phages were 2811, 311F and 05F, totalling 6 phages in the final cocktail. There was no lytic activity of the *Klebsiella* phages against the *E. coli* clinical isolates or vice-versa based on DST [Supplementary Data Table S3 and S4]. The phage genomes were analysed to ensure that they did not carry genes known to allow a lysogenic lifestyle and did not contain any genes encoding for known toxins. A summary of the characteristics of the final six phages are shown in Figure 2.

### Use of virulence index score demonstrates synergy within phage combinations

#### Analysis of the combinations of ESBL *E. coli* phages using virulence index scores

Phages UP17, JK08 and 113 used in various combinations of doublets, triplets and also in the final ESBL six-phage cocktail were tested using the planktonic killing assays. Using the quantitative virulence index scores, all data was compared (Table 2). Data was compared on two scales; the macroscale to analyse only the number of clinical isolates within each virulence index category and the microscale to analyse individual clinical isolate virulence index scores for each phage combination.

Based on the virulence index data from the three individual phages (UP17, JK08 and 113), 13/19 (68%) of *E. coli* isolates should be targeted, however only 12/19 (63%) were (Table 1). There was an unexpected reduction in the number of isolates killed by the triplet phage combination (63%) when compared with the final six phage combination (53%) (Table 1).

**Table 1.** The virulence index scores of each ESBL-producing *E. coli* clinical isolates against each *E. coli* phage combination summarised into categories. The rows represent the categories, high = virulence index score > 0.5, medium = virulence index score 0.2 - 0.5, low = virulence index score ≥ 0.001 - < 0.2, none = 0. Effective is a combination of the high and medium categories, this defines effective killing by the phage combination and its clinical isolate coverage. The columns represent the various phage combinations from left to right commencing with the doublets (UP17&JK08, UP17&113 and JK08&113), triplet (UP17, JK08 & 113) and final (UP17, JK08, 113, 2811, 311F and 05F).

|  | UP17&JK08 | UP17&113 | JK08&113 | Triplet | Final |
| --- | --- | --- | --- | --- | --- |
| High<br>> 0.5 | 10 | 6 | 5 | 8 | 8 |
| Medium<br>0.2 - 0.5 | 2 | 0 | 5 | 4 | 2 |
| Low<br>> 0.001 - < 0.2 | 5 | 3 | 5 | 4 | 6 |
| None<br>0 | 2 | 10 | 4 | 3 | 3 |
| Effective<br>(High & Medium) | 12/19<br>63% | 6/19<br>32% | 10/19<br>53% | 12/19<br>63% | 10/19<br>53% |

When comparing virulence index scores, there were no substantial differences between the triplet cocktail (UP17, JK08 & 113) and the final six phage cocktail for the majority of the individual clinical isolates (Table 2). However, the virulence index identified inhibitory combinations. For example, when KR2729 was treated with phage 113 alone a high virulence index score of 0.64 is obtained (Table 2). But when used in combination with phage JK08 (JK08 & 113), its virulence index score dropped to almost zero (0.05) (Table 2). When all three phages were used in combination, the high virulence index score is restored to 0.92, which could be due to phage UP17 alone (Table 2). This effect is only noted where phage 113 is the only phage to have a high virulence index score, but with no noticeable effect from phage JK08 (Table 2). The effect was not noted in combinations where both phages JK08 and 113 had medium or high virulence index scores. This was exemplified by clinical isolates MH10 & MH14 (Table 2).

**Table 2.**
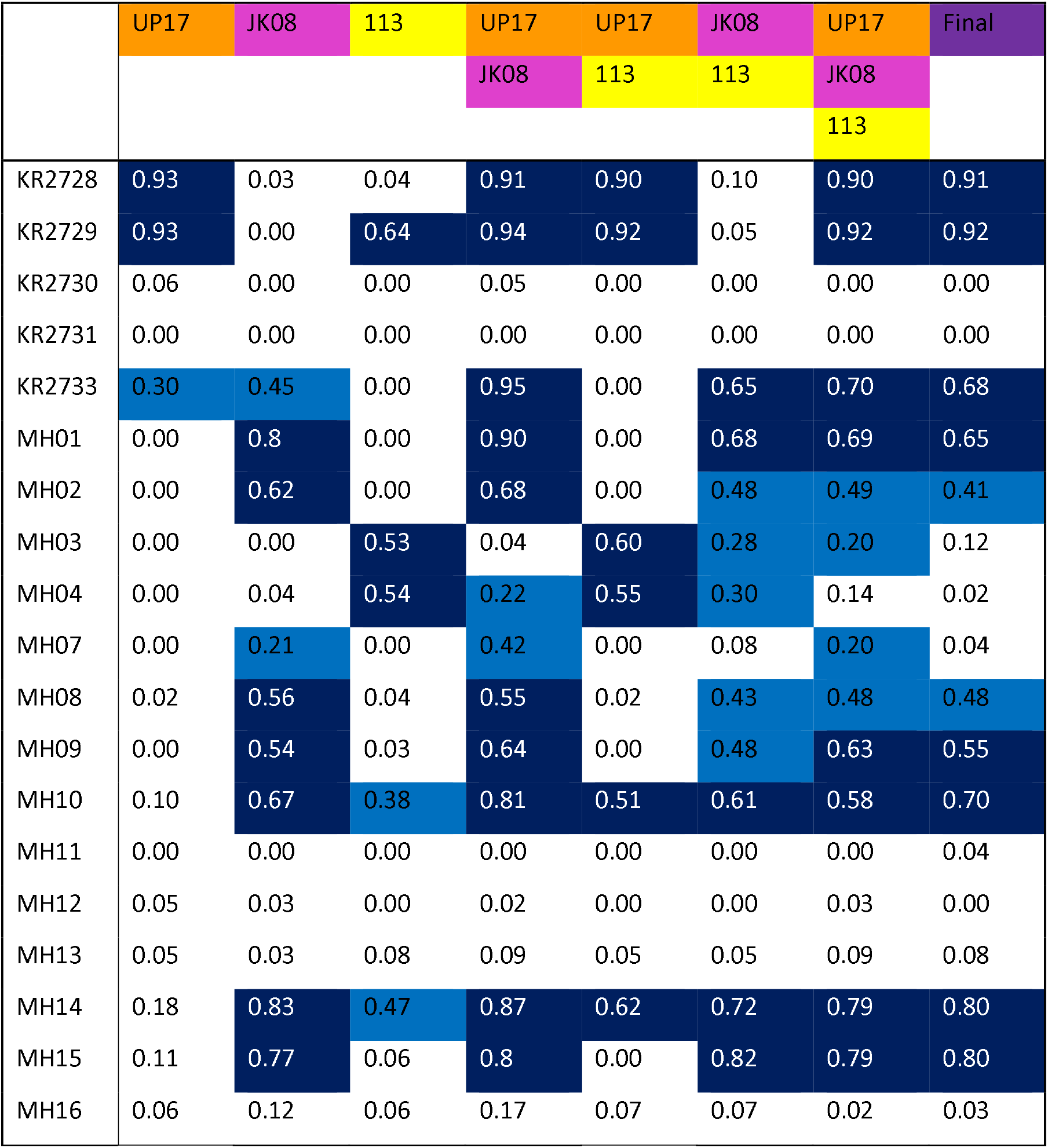
The virulence index scores of individual phage and various phage combinations across all the 19 ESBL-producing *E. coli* clinical isolates. Numbers highlighted in Dark Blue represent high virulence index scores and those highlighted in Light Blue represent medium virulence index scores. The rows represent each of the ESBL-producing *E. coli* clinical isolates used within this study. The top column represents the phage starting with individual phage on the left to progressing across the various combinations. The phages are highlighted with different colours: UP17 (Orange), JK08 (Pink), 113 (Yellow), Final (Purple). Final = all six final phage (UP17, JK08, 113, 2811, 311F and 05F). All values represent the mean generated from triplicate experimental data. A full diagrammatic representation of this data can be seen in Supplementary Data Figure S7.

Conversely synergistic interactions were also observed. Treating KR2733 with phage UP17 or JK08 results in virulence index scores of 0.3 and 0.45, respectively. However, when used in combination the virulence index increases to 0.95 (Table 2). A similar pattern can be seen for clinical isolates, MH01, MH10 and MH07 (Table 2) with this phage combination.

#### Analysis of the combinations of ESBL *Klebsiella* phages using virulence index scores

The same selection process was carried out for comparison of *Klebsiella* phages. The effectiveness of different combinations of phages 2811, 311F and 05F was compared using the virulence index scores to assess the efficacy (Table 3). The most effective doublet combination was 311F & 05F, which targets 53% of isolates. The addition of a further phage had a detrimental effect, reducing the number of isolates killed to 37% (Table3).

**Table 3.** The virulence index scores of each ESBL-producing *Klebsiella* clinical isolates against each *Klebsiella* phage combination summarised into categories. The rows represent the categories, high = virulence index score > 0.5, medium = virulence index score 0.2 - 0.5, low = virulence index score ≥ 0.001 - < 0.2, none = 0. Effective = a combination of the high and medium categories, this defines effective killing by the phage combination and its clinical isolate coverage. The columns represent the various phage combinations from left to right commencing with the doublets (2811&311F, 2811&05F and 311F&05F), triplet (2811, 311F & 05F) and final (UP17, JK08, 113, 2811, 311F and 05F).

|  | 2811&311F | 2811&05F | 311F&05F | Triplet | Final |
| --- | --- | --- | --- | --- | --- |
| High<br>> 0.5 | 3 | 2 | 1 | 3 | 2 |
| Medium<br>≥ 0.2 - 0.5 | 1 | 4 | 9 | 4 | 11 |
| Low<br>> 0.001 – < 0.2 | 12 | 12 | 9 | 12 | 6 |
| None<br>0 | 3 | 1 | 0 | 0 | 0 |
| Effective (High<br>& Medium) | 4/19<br>(21%) | 6/19<br>(32%) | 10/19<br>(53%) | 7/19<br>(37%) | 13/19<br>(68%) |

Seven clinical isolates are targeted by the triplet cocktail compared to the six isolates covered based on the individual phage data (Table 4). The additional clinical isolate targeted by the triplet, KR398, showed a virulence index score (0.22) just above the threshold (Table 4). This suggests that for the *Klebsiella* phages, the killing seen with the individual phage translates directly to the triplet combination of phages. Additionally, the virulence index scores of the individual phage and of the triplet suggesting no synergy or competitive inhibition across all the clinical isolates. For example, clinical isolate KR438, phage 2811 only (0.78), triplet (0.73) or clinical isolate MH05 phage 05F only (0.32), triplet (0.33) (Table 4).

**Table 4.**
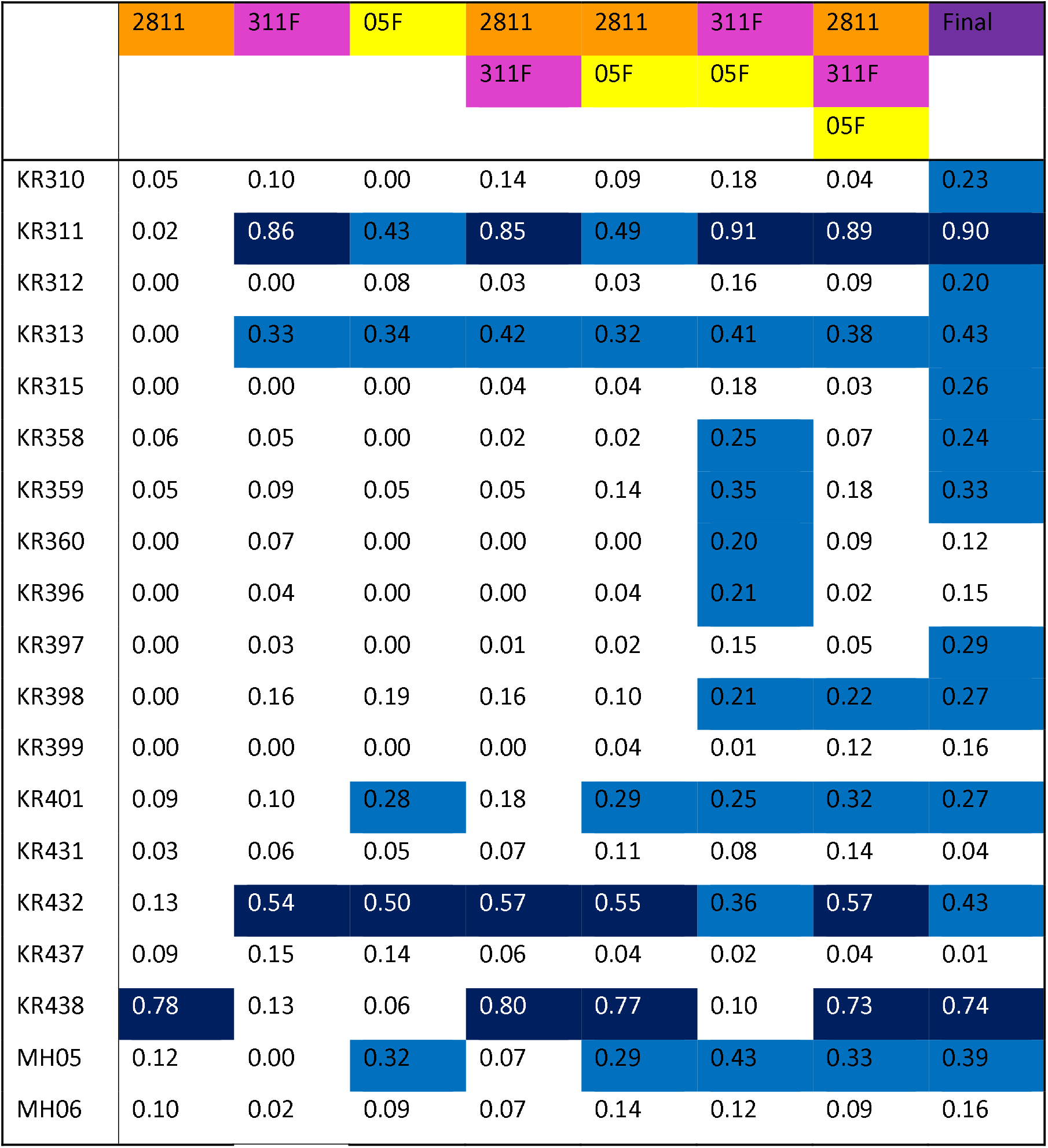
The virulence index scores of individual phage and various phage combinations across all the 19 ESBL-producing *Klebsiella* clinical isolates. Numbers highlighted in Dark Blue represent high virulence index scores and those highlighted in Light Blue represent medium virulence index scores. The rows represent each of the ESBL *Klebsiella* clinical isolates used within this study. The top column represents the phage starting with individual phage on the left to progressing across the various combinations. The phages are highlighted with different colours: 2811 (Orange), 311F (Pink), 05F (Yellow), Final (Purple). Final = all six final phage (UP17, JK08, 113, 2811, 311F and 05F). All values represent the mean generated from triplicate experimental data. A full diagrammatic representation of this data can be seen in Supplementary Data Figure S7.

|  | 2811 | 311F | 05F | 2811 | 2811 | 311F | 2811 | Final |
| --- | --- | --- | --- | --- | --- | --- | --- | --- |
|  |  |  |  | 311F | 05F | 05F | 311F |  |
|  |  |  |  |  |  |  | 05F |  |
| KR310 | 0.05 | 0.10 | 0.00 | 0.14 | 0.09 | 0.18 | 0.04 | 0.23 |
| KR311 | 0.02 | 0.86 | 0.43 | 0.85 | 0.49 | 0.91 | 0.89 | 0.90 |
| KR312 | 0.00 | 0.00 | 0.08 | 0.03 | 0.03 | 0.16 | 0.09 | 0.20 |
| KR313 | 0.00 | 0.33 | 0.34 | 0.42 | 0.32 | 0.41 | 0.38 | 0.43 |
| KR315 | 0.00 | 0.00 | 0.00 | 0.04 | 0.04 | 0.18 | 0.03 | 0.26 |
| KR358 | 0.06 | 0.05 | 0.00 | 0.02 | 0.02 | 0.25 | 0.07 | 0.24 |
| KR359 | 0.05 | 0.09 | 0.05 | 0.05 | 0.14 | 0.35 | 0.18 | 0.33 |
| KR360 | 0.00 | 0.07 | 0.00 | 0.00 | 0.00 | 0.20 | 0.09 | 0.12 |
| KR396 | 0.00 | 0.04 | 0.00 | 0.00 | 0.04 | 0.21 | 0.02 | 0.15 |
| KR397 | 0.00 | 0.03 | 0.00 | 0.01 | 0.02 | 0.15 | 0.05 | 0.29 |
| KR398 | 0.00 | 0.16 | 0.19 | 0.16 | 0.10 | 0.21 | 0.22 | 0.27 |
| KR399 | 0.00 | 0.00 | 0.00 | 0.00 | 0.04 | 0.01 | 0.12 | 0.16 |
| KR401 | 0.09 | 0.10 | 0.28 | 0.18 | 0.29 | 0.25 | 0.32 | 0.27 |
| KR431 | 0.03 | 0.06 | 0.05 | 0.07 | 0.11 | 0.08 | 0.14 | 0.04 |
| KR432 | 0.13 | 0.54 | 0.50 | 0.57 | 0.55 | 0.36 | 0.57 | 0.43 |
| KR437 | 0.09 | 0.15 | 0.14 | 0.06 | 0.04 | 0.02 | 0.04 | 0.01 |
| KR438 | 0.78 | 0.13 | 0.06 | 0.80 | 0.77 | 0.10 | 0.73 | 0.74 |
| MH05 | 0.12 | 0.00 | 0.32 | 0.07 | 0.29 | 0.43 | 0.33 | 0.39 |
| MH06 | 0.10 | 0.02 | 0.09 | 0.07 | 0.14 | 0.12 | 0.09 | 0.16 |

Analysis of the doublet (311F & 05F) showed unexpected synergistic combination. For five clinical isolates (KR358, KR359, KR360, KR396 and KR398), individually phages 311F and 05F had an almost negligible effect, but when combined (311F & 05F) they demonstrated medium virulence index scores for all strains (Table 4). For the triplet cocktail (2811, 311F and 05F), five clinical isolates (KR310, KR312, KR315, KR358, KR359, KR397) again had negligible virulence index scores (Table 4). But when exposed to the final cocktail (UP17, JK08, 113, 2811, 311F and 05F), all five clinical isolates had a medium virulence index scores (Table 4). This demonstrated a further unexpected synergy when added with the ESBL *E. coli* phages.

### Effectiveness of the Final ESBL Phage Cocktail

The final ESBL cocktail was effective against 23/38 clinical isolates (61%) based on the virulence index data (any clinical isolates with a medium or high virulence index score > 0.2). The final ESBL cocktail was then tested in a 24-hour biofilm assay, to test the cocktail in a bacterial virulence model.

The final ESBL cocktail was most effective against the *E. coli* clinical isolates. There was a significant (*p* < 0.05) decrease in bacterial cell viability in 11 (58%) and 13 (68%) isolates after 4 and 24 hours of resazurin incubation respectively (Figure 3).

**Figure 3.**
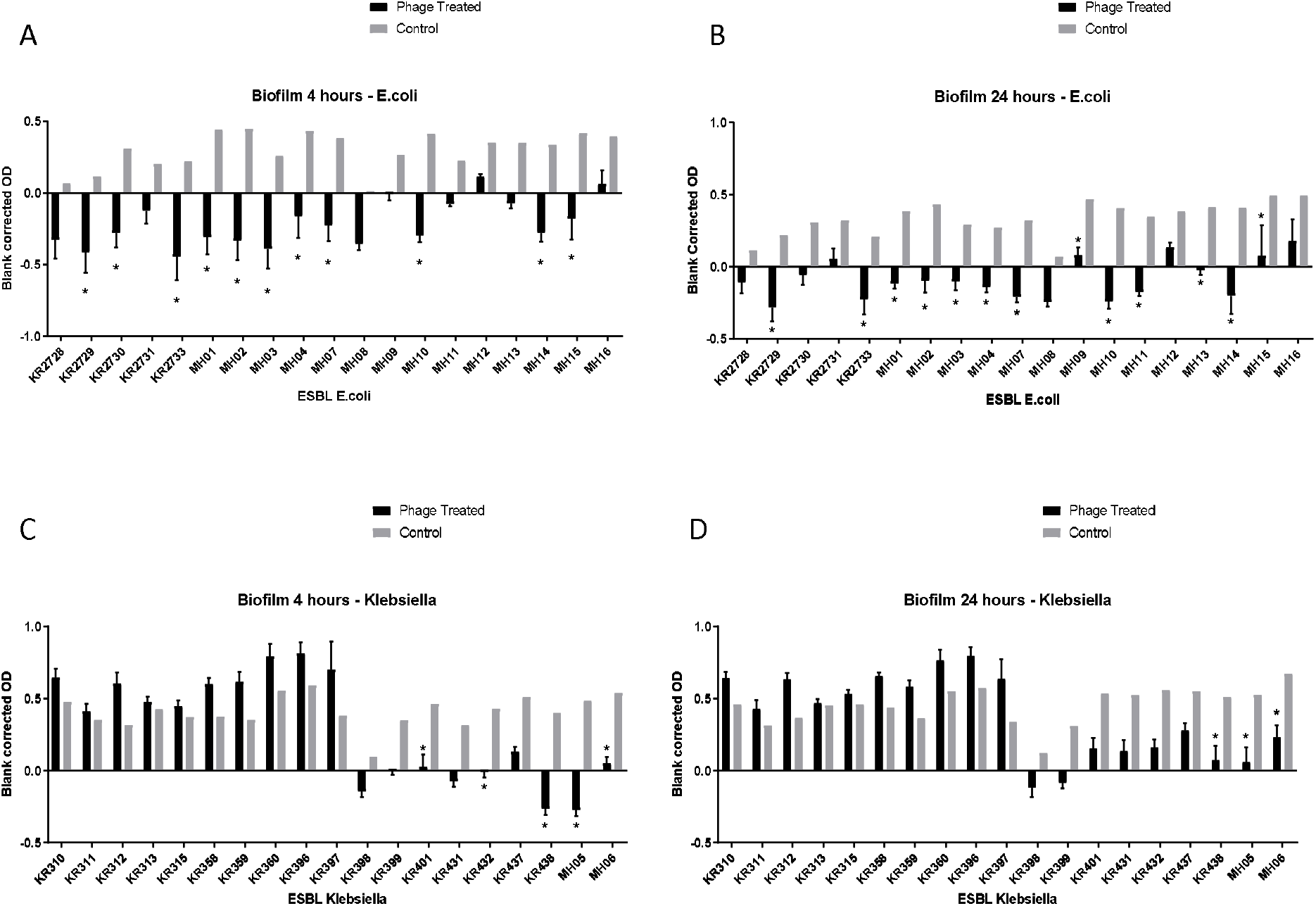
Graphical representation of the biofilm assay data – resazurin cell viability-based model on a 96 well plate. Clinical isolates were grown for 24 hours on a flat bottom 96 well plate, then incubated for an additional 24 hours with either LB (control) or the final phage cocktail (phage treated). Resazurin was then added, OD readings were taken at 4 hours and 24 hours post adding resazurin. Each ESBL-producing clinical isolate has two bars: the black bar represents the phage treated blank corrected OD and the grey bar represents the control blank corrected OD. OD taken at A_595_, experiments repeated in triplicate for all clinical isolates, columns represents mean with standard error of the mean. *=significance difference between those treated with final phage cocktail and the control, *p* <0.05. The top left graph (A) depicts the all ESBL producing E.coli clinical isolates 4 hours post incubation with resazurin, top right (B) depicts ESBL producing E.coli clinical isolates 24 hours post incubation with resazurin, the bottom left (C) depicts all ESBL-producing *Klebsiella* clinical isolates 4 hours post incubation with resazurin and bottom right (D) depicts all ESBL-producing *Klebsiella* clinical isolates 24 hours post incubation with resazurin.

For *Klebsiella*, at 4 hours the cocktail only killed 5/19 (26%) of isolates and at 24 hours 3/19 (16%) (Figure 3). This in stark contrast to the high clinical isolate killing observed by the planktonic killing assay of 13/19 (68%) (Table 3). An example of the disparity of results between the two tests is clinical isolate, KR311. It had the highest virulence index score of 0.9 (Table 4), when using the final ESBL cocktail in the planktonic killing assay but had no significant (*p* < 0.05) decrease in bacterial cell viability (Figure 3). However, the second highest virulence index score of 0.74 on isolate KR438 (Table 4) correlated with a significant (*p* < 0.05) reduction in the biofilm assay (Figure 3).

### Genomic analysis of the ESBL-producing clinical isolates

Core genome SNP analysis was used to compare the clinical isolates (Figure 4). Ten different ST types of *E. coli* were identified, with the cocktail being able to target ten strains across three ST types. The cocktail could target 5/6 of the ST131 clinical isolates, which is the most prevalent multidrug resistant uropathogen (Johnson et al., 2010; Kudinha et al., 2013). The core-genome SNP analysis of *Klebsiella*, clearly separated the isolates into two different species (Figure 4). Three isolates were *Klebsiella oxytoca* and the remainder *Klebsiella pneumoniae*. There was a broad diversity of ST types present with 12 different types detected. There are representatives of the global endemic carbapenem-resistant associated ST258 as well as high risk AMR type ST147 (Bowers et al., 2015; Dhar et al., 2016; Peirano et al., 2020). The cocktail of phages was able to target a broad diversity of ST types across the three different bacterial species.

**Figure 3.**
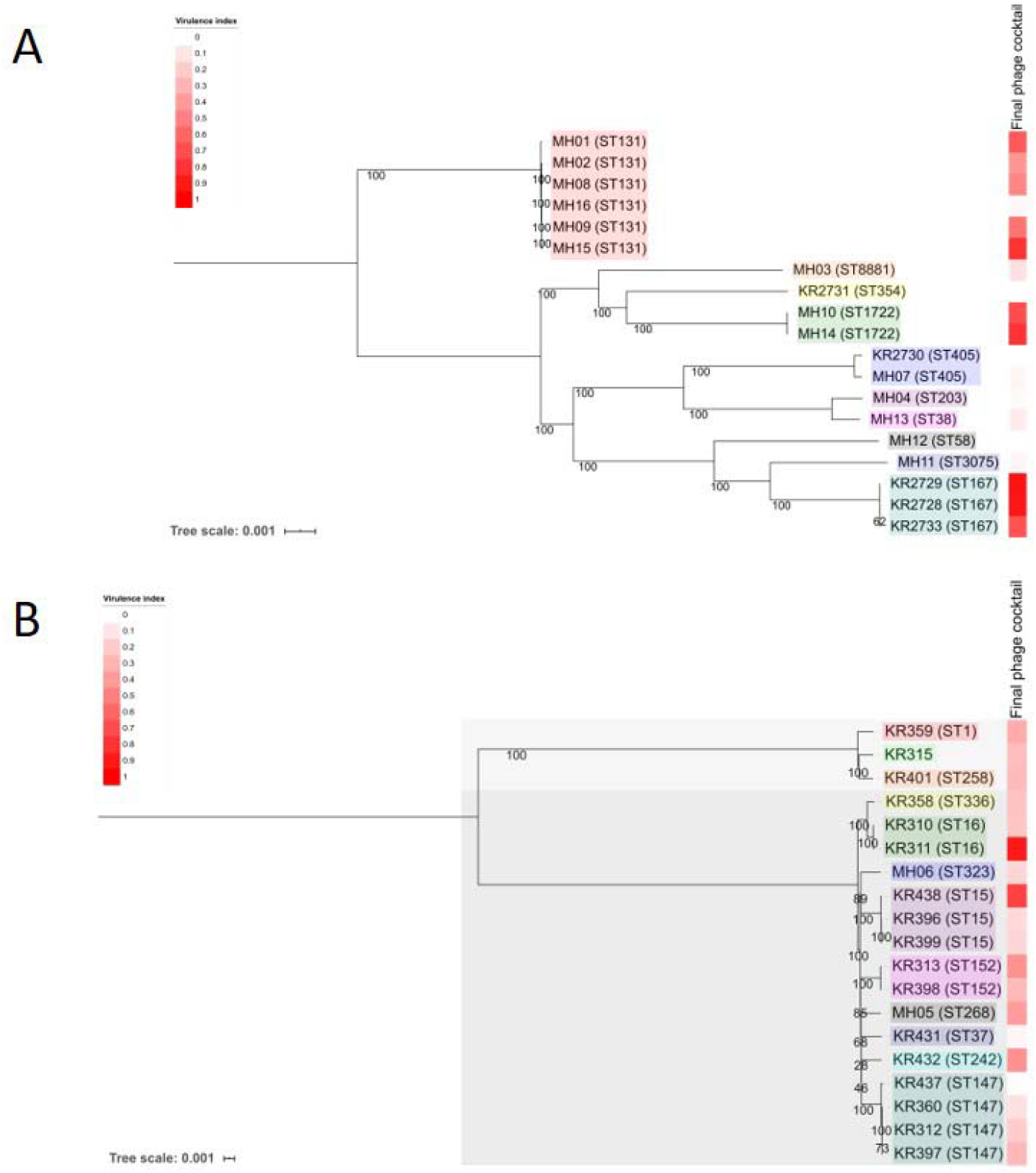
Phylogenetic analysis of the 38 ESBL-producing clinical isolates used in this study. Panel A shows ESBL-producing *E. coli* and Panel B shows ESBL-producing *Klebsiella* spp. Trees were produced using MEGA7 to assess the core genome SNPs. Core-genome SNP analysis revealed that there were two species of *Klebsiella*. KR315, KR359 and KR401 are *Klebsiella* oxytoca and all others are *Klebsiella* pneumoniae. Each clinical isolate name is followed by its MLST (Achtman – *E. coli*) – please note KR315 was unable to be assigned. Tree scale noted and bootstrap values are labelled on branches. Coloured boxes within each tree represent groups of sequence types. The heat map on each tree represents the virulence index score assigned to the final phage cocktail for each strain.

Figure 5 allows an overview of the planktonic killing assay virulence index scores taking into consideration all phage combinations including individual, doublets, triplets and the six-phage final cocktail that were used during this work. It also includes combinations using phages that were screened but not selected as the final six phages.

**Figure 4.**
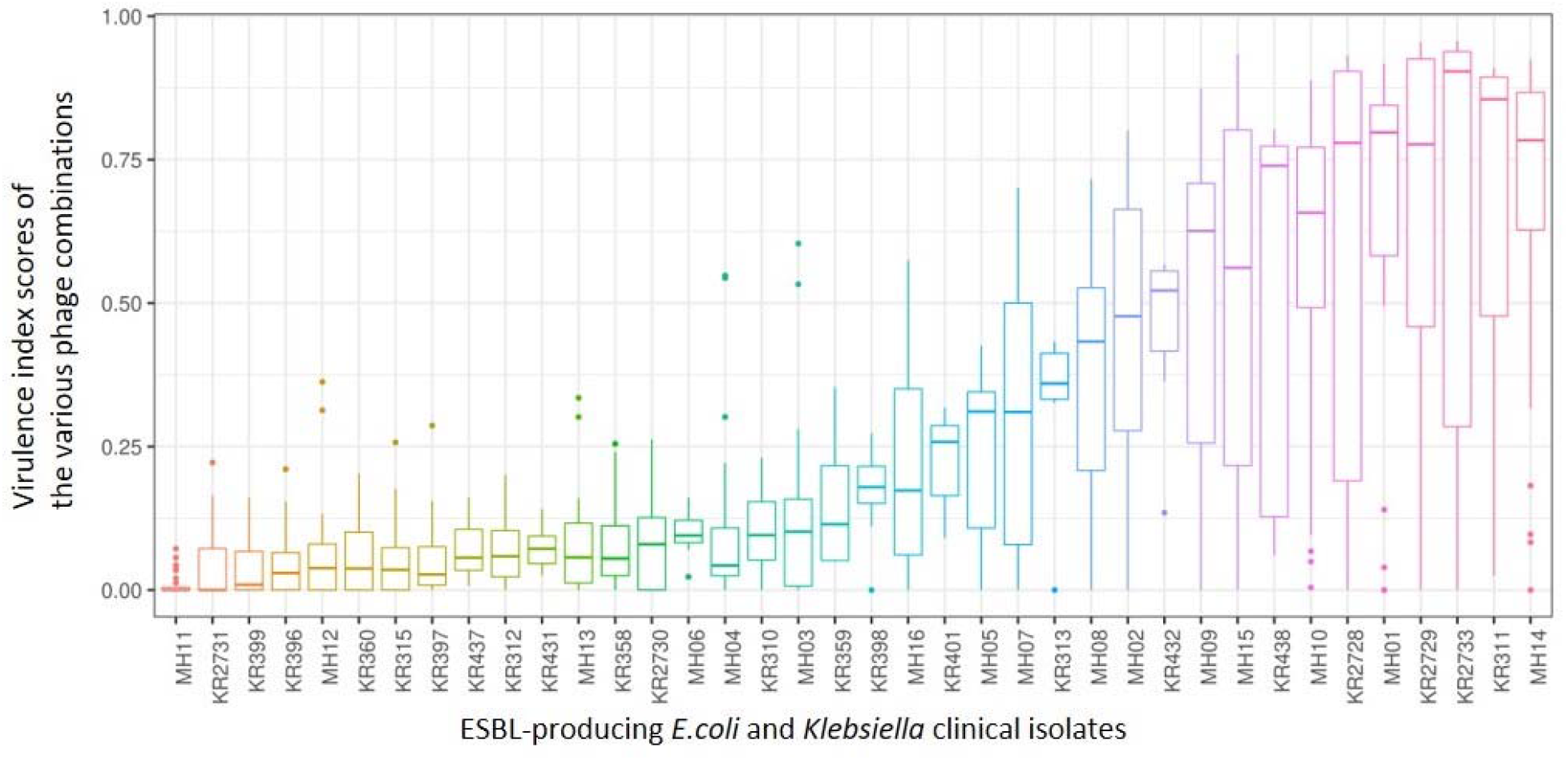
A box and whisker plot depicting the range of killing assay virulence index scores from all phage combination explored during this work for each of the 38 ESBL-producing clinical isolates. From left to right shows the ESBL-producing clinical isolates that are most resistant to the combinations attempted to those that are most sensitive. Please note not all combinations were completed in triplicate for those combinations that were not part of the final ESBL cocktail (all six phages).

When comparing this data with the two phylogenetic trees (Figure 4) of all 38 clinical isolates, there is no clear pattern of genomic similarities to phage susceptibility. The most sensitive *E. coli* clinical isolates (MH14, KR2733, KR2729, MH01, KR2728) are spread across three different clades. In contrast the most resistant clinical isolates were spread across five different clades (MH11, KR2731, MH12, MH13 and KR2730). With regards to *Klebsiella*, the most sensitive strains were spread across five clades (KR311, KR438, KR432, KR313, MH05). The most resistant strains were distributed across three different clades (KR399, KR396, KR360, KR315, KR397).

## Discussion

Antimicrobial resistance is an urgent issue that needs to be addressed. Phage therapy could be part of the solution. This work focuses on the development of an effective phage cocktail in response to this need. The aim of this work was to assess phage selection methods to streamline the development of a phage cocktail. This was achieved using an example of a phage cocktail against ESBL-producing clinical isolates of *E. coli* and *Klebsiella*.

DST is commonly used in the literature to assess the host range of phage (Hyman, 2019; Montso et al., 2019; Sybesma et al., 2016). The data demonstrated that the DST overestimates the host range or clinical isolate coverage of the individual phage by approximately 50% compared with the EOP and the planktonic killing assay (Figure 2). The discrepancy between DST and EOP is in keeping with previous publications relating to Enterobacteriaceae species (Manohar et al., 2019; Mirzaei and Nilsson, 2015). It is considered to be due to other mechanisms of killing noted with DST, such as ‘lysis from without,’ that are not a result of phage replication and lysis of the bacterial cell (Hyman and Abedon, 2010). But when comparing EOP with planktonic killing assay data, there is less disparity in the numbers of clinical isolates and specific individual clinical isolates covered.

Use of the virulence index score for analysis across a large dataset allowed direct comparison of individual phages and phage combinations, which would be a powerful tool for cocktail design. Overall, based on the dataset generated by this work, there is not a clear formula for the expected outcome when combining phages. This is due to either synergy or inhibition, which cannot be easily predicted. The ability of the virulence index to detect these interactions is a clear advantage over the use of DST or EOP as a selection method. The concepts of viral interference and augmentation have previously been discussed in the literature (Casey et al., 2018). Synergistic enhancement could be due to an effect on one or more of the three properties: 1) rate of infection, 2) production of progeny or 3) the time window between infection and progeny release (Schmerer et al., 2014). Therefore, synergy of phage infection is an additional advantage for the creation of a successful phage cocktail (Schmerer et al., 2014). The planktonic killing assay method alongside analysis using the virulence index score could make this a realistic research aim during future cocktail design. UP17 was interesting in that it also appeared to be resistant to interference from the other phages within the cocktail. This is shown with Table 2, where UP17 had a high virulence score against a particular clinical isolate (KR2728 & KR2729) this score is maintained throughout all the other phage combinations with UP17 (UP17&JK08, UP17&113, UP17&JK08&113, all 6).

Overall, it would be worth investigating further, why UP17 is resistant to interference from the other phages as well as to why its effectiveness increases when combined with JK08. In addition, why JK08 and 113 had an antagonist relationship. This could be due to the phages having similar receptor sites and one being more likely to lead to an abortive infection, or superinfection resulting in an unsuccessful infection for both (Abedon, 2015). Answering all of these questions, may help determine effective future cocktail design. The strength of this work is the use of the virulence index score to be able to support the combination of phages together in a cocktail by providing clear evidence of synergy. This synergy would not be apparent from other commonly used selection methods such as DST and EOP. In addition, this method also outperforms the previous planktonic killing assay methods, with the use of time course measurements in a 96-well plate format, as it allows high throughput of a large number of individual phage/phage combinations and clinical isolate panels.

The data suggests that other factors may come into play for *Klebsiella clinical* isolates. When comparing the phage virulence assays of the biofilm assay and planktonic killing assay, there appears to be no correlation for *Klebsiella* clinical isolates. For example, when using the final ESBL cocktail the clinical isolate KR311 has the highest virulence index score of 0.9 in the planktonic killing assay (Table 4) and yet there was no significant (*p* < 0.05) decrease in the biofilm assay (Figure 3). In contrast, the second highest virulence index score of 0.74 against KR438 correlated with a significant (*p* < 0.05) reduction in cell viability in the biofilm assay (Table 4, Figure 3). Overall, when assessing the clinical isolates that demonstrated a significant (*p* < 0.05) reduction in cell viability for at least one of the timepoints during the biofilm assay, there appears to be no correlation with the planktonic killing assay virulence index. This is a disappointing result, at least for this biofilm model, as the ideal case would be for the high-throughput method to determine virulence as the planktonic killing assay to translate to effectiveness in an *in vitro* bacterial model of virulence. The final cocktail covered 13 isolates (68%) in the planktonic killing assay (Table 3) in comparison to 5/19 (26%) clinical isolates at 4 hours and 3/19 (16%) clinical isolates at 24 hours within the biofilm assay (Figure 3). It has been demonstrated in the literature that the use of phage can cause a significant reduction in biofilm production in *Klebsiella* spp. (Tabassum et al., 2018; Taha et al., 2018). This is in conflict to data demonstrated in this work. The results with *E. coli* were more promising as the clinical isolate coverage was similar 10/19 (53%) for planktonic killing assay and in the biofilm assay 11/19 (58%) at 4 hours and 13/19 (68%) at 24 hours. The overall aim of this work was to develop a phage cocktail that was effective against 90% of the ESBL-producing clinical isolate collection. This was not achieved, and the result may influence the clinical application of the final cocktail. The data presented here demonstrated it was highly effective against the global prominent AMR UTI-associated *E. coli* ST131 isolates with 5/6 (83%) isolates killed. The data in this paper would need to be reconciled with prevalence data of the different sequence types within the general population to be able to draw conclusions with regards the true clinical application.

In this study, we also performed basic bacterial genetic analysis identifying those that were most resistant to phage infection and those that were most sensitive. This highlighted clinical isolates that were on the same clade on the phylogenetic tree (Figure 4) but have polar opposite phage sensitivity. An example, *Klebsiella* KR396 and KR399 are both resistance isolates against sensitive isolate *Klebsiella* KR438 (Figure 4). Further genetic analysis of those with polar opposite phage sensitivity could provide further insight into mechanisms of resistance. Further genetic analysis could provide an opportunity to assess the individual clinical isolates susceptibility across three different screening methods and biofilm assay to see if there were any markers that predicted the outcome. These markers could help in the design of cocktails. In the future, a more detailed genomic analysis of the clinical isolates will be reported.

This paper is intended to outline the selection of phages for a final cocktail formulation. There are two methods that can be considered for use (Figure 6). As per the method used in this study begin by filtering the potential phage candidates via DST, then further screen with EOP. Then proceed to use the EOP data to assess for coverage of the selected bacterial isolate panel followed by testing the combinations in a high throughput planktonic killing assay. The alternative to consider is to use the DST data and then immediately proceed to a planktonic killing assay using individual phage. Then use the single phage planktonic killing assay data to select phage for the cocktail combination.

**Figure 5.**
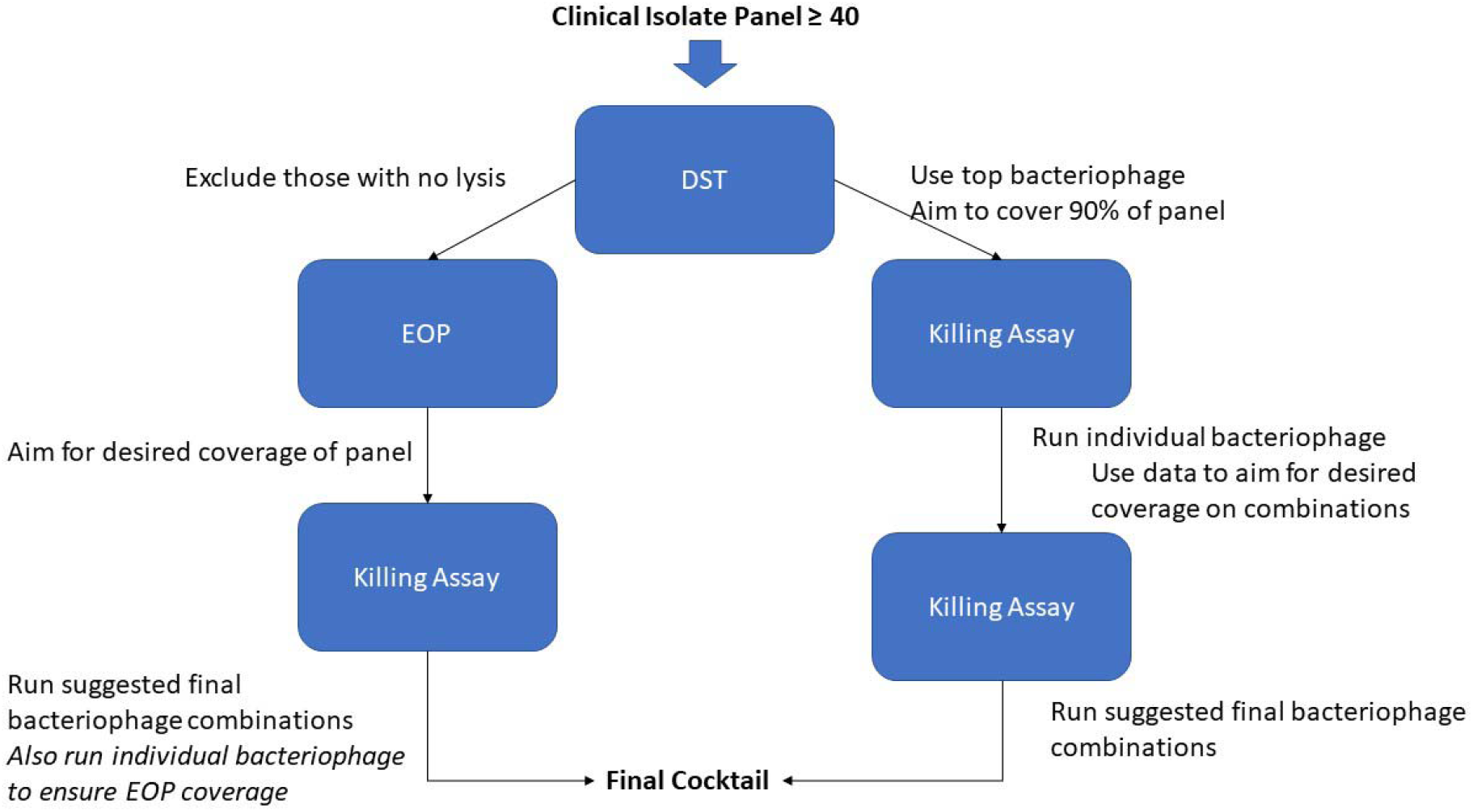
depicts the two suggested processes to screen a phage library against a clinical isolate collection to optimise development of a therapeutic phage cocktail.

In conclusion, DST and EOP are not as useful as the planktonic assay as selection methods for designing phage cocktails. This is due to the inability of the DST and EOP to identify beneficial synergy as well as avoid inhibition.

## Acknowledgements

Thanks to Mohammed Imam and Wafaa Alrashidi for access to their phages used during initial screening, but not used in final ESBL cocktail. Thanks to University of Leicester and Marialuisa Crosatti for access to the Kumar Rajakumar bacterial strain collection. Thanks to Shaun Livesey from University Hospitals of Leicester for support with completing the ESBL producing clinical isolate collection. Thanks to Lucy Gannon and Christian Harrison for technical support. Thanks to Natalie Allcock from University of Leicester for producing Transmission Electron Microscopy Images for UP17, 113, 2811, 311F and 05F. Thanks to Horst Neve at MRI, Kiel, Germany for producing Transmission Electron Microscopy Images for JK08. Thanks to Gurinder Vinner for support with laboratory skills and experiment planning.

This publication made use of the PubMLST website (https://pubmlst.org/) developed by Keith Jolley (Jolley & Maiden 2010, BMC Bioinformatics, 11:595) and sited at the University of Oxford. The development of that website was funded by the Wellcome Trust.

## Funding

MH is funded by a National Institute for Health Research (NIHR) Academic Clinical Fellowship for this research project. The views expressed in this publication are those of the author(s) and not necessarily those of the NIHR or the Department of Health and Social Care. BA was funded by the Ministry of Education in Saudi Arabia as a PhD sponsorship to BA (KSU/1480/E). JM and DvS were partly supported by a grant generously provided by the Bill & Melinda Gates Foundation (Ref. No. OPP1150567). JM is also supported by a Starting Investigator Research Grant (SIRG) (Ref. No. 15/SIRG/3430) funded by Science Foundation Ireland (SFI). DvS supported by a Principal Investigator award (Ref. No. 13/IA/1953) through SFI. FH is funded by the Biotechnology and Biological Sciences Research Council (BBSRC) Midlands Integrative Biosciences Training Partnership for this project. AM was funded by Natural Environment Research Council grants (NE/N019881/1 & NE/N003241/1) and MRC CLIMB (MR/L015080/1).

## Author Contributions Statement

MH and MC designed the experiments. MH, JN, JM, DvS, JK, BA, MA provided the phage used in this study. MH performed the DST, planktonic killing assays and biofilm assays. MH & FH performed the EOP assays. JK, NB and AT prepared phage genomic DNA for sequencing. FH and AM performed the bioinformatic analysis of the phage and bacterial genomes. MH, FH, DS, NB analysed the data. MH interpreted the results. MH drafted the manuscript. FH, JN, JM, DS, AT, AM, EG and MC edited the manuscript.

## Notes

### Competing Interest Statement

The authors have declared no competing interest.

## References

Abedon, S. T. (2015). Bacteriophage secondary infection. Virol. Sin. 30, 3–10. doi:10.1007/s12250-014-3547-2.

Bankevich, A., Nurk, S., Antipov, D., Gurevich, A. A., Dvorkin, M., Kulikov, A. S., et al. (2012). SPAdes: A new genome assembly algorithm and its applications to single-cell sequencing. J. Comput. Biol. 19, 455–477. doi:10.1089/cmb.2012.0021.

Bolger, A. M., Lohse, M., and Usadel, B. (2014). Trimmomatic: A flexible trimmer for Illumina sequence data. Bioinformatics 30, 2114–2120. doi:10.1093/bioinformatics/btu170.

Bowers, J. R., Kitchel, B., Driebe, E. M., MacCannell, D. R., Roe, C., Lemmer, D., et al. (2015). Genomic Analysis of the Emergence and Rapid Global Dissemination of the Clonal Group 258 Klebsiella pneumoniae Pandemic. PLoS One 10, e0133727. doi:10.1371/journal.pone.0133727.

Bush, K. (2018). Past and present perspectives on β-lactamases. Antimicrob. Agents Chemother. 62. doi:10.1128/AAC.01076-18.

Casey, E., van Sinderen, D., and Mahony, J. (2018). In vitro characteristics of phages to guide ‘real life’ phage therapy suitability. Viruses 10. doi:10.3390/v10040163.

Chevreux, B., Wetter, T., and Suhai, S. (1999). Genome Sequence Assembly Using Trace Signals and Additional Sequence Information. Comput. Sci. Biol. Proc. Ger. Conf. Bioinforma.’99, GCB, Hann. Ger., 45–56. doi:10.1.1.23/7465.

Dhar, S., Martin, E. T., Lephart, P. R., McRoberts, J. P., Chopra, T., Burger, T. T., et al. (2016). Risk factors and outcomes for carbapenem-resistant Klebsiella pneumoniae isolation, stratified by its multilocus sequence typing: ST258 versus non-ST258. Open Forum Infect. Dis. 3. doi:10.1093/ofid/ofv213.

Feldgarden, M., Brover, V., Haft, D. H., Prasad, A. B., Slotta, D. J., Tolstoy, I., et al. (2019). Validating the AMRFINder tool and resistance gene database by using antimicrobial resistance genotype-phenotype correlations in a collection of isolates. Antimicrob. Agents Chemother. 63. doi:10.1128/AAC.00483-19.

Garneau, J. R., Depardieu, F., Fortier, L. C., Bikard, D., and Monot, M. (2017). PhageTerm: A tool for fast and accurate determination of phage termini and packaging mechanism using next-generation sequencing data. Sci. Rep. 7, 1–10. doi:10.1038/s41598-017-07910-5.

Gurevich, A., Saveliev, V., Vyahhi, N., and Tesler, G. (2013). QUAST: Quality assessment tool for genome assemblies. Bioinformatics 29, 1072–1075. doi:10.1093/bioinformatics/btt086.

Hyman, P. (2019). Phages for phage therapy: Isolation, characterization, and host range breadth. Pharmaceuticals 12, 35. doi:10.3390/ph12010035.

Johnson, J. R., Johnston, B., Clabots, C., Kuskowski, M. A., and Castanheira, M. (2010). Escherichia coli sequence type ST131 as the major cause of serious multidrug-resistant E. coli infections in the United States. Clin. Infect. Dis. 51, 286–294. doi:10.1086/653932.

Kropinski, A. M., Mazzocco, A., Waddell, T. E., Lingohr, E., and Johnson, R. P. (2009). Chapter 7: Enumeration of Bacteriophages by Double Agar Overlay Plaque Assay. Methods Mol. Biol. 501, 69–76. doi:10.1007/978-1-60327-164-6_7.

Kudinha, T., Johnson, J. R., Andrew, S. D., Kong, F., Anderson, P., and Gilbert, G. L. (2013). Escherichia coli sequence type 131 as a prominent cause of antibiotic resistance among urinary Escherichia coli isolates from reproductive-age women. J. Clin. Microbiol. 51, 3270–3276. doi:10.1128/JCM.01315-13.

Kumar, S., Stecher, G., Li, M., Knyaz, C., and Tamura, K. (2018). MEGA X: Molecular Evolutionary Genetics Analysis across Computing Platforms. Mol. Biol. Evol. 35, 1547–1549. doi:10.1093/molbev/msy096.

Kutter, E. (2009). Chapter 14: Phage Host Range and Efficiency of Plating. Methods Mol. Biol. 501, 141–149. doi:10.1007/978-1-60327-164-6_14.

Letunic, I., and Bork, P. (2007). Interactive Tree Of Life (iTOL): an online tool for phylogenetic tree display and annotation. Bioinformatics 23, 127–8. doi:10.1093/bioinformatics/btl529.

Li, D., Liu, C. M., Luo, R., Sadakane, K., and Lam, T. W. (2015). MEGAHIT: An ultra-fast single-node solution for large and complex metagenomics assembly via succinct de Bruijn graph. Bioinformatics 31, 1674–1676. doi:10.1093/bioinformatics/btv033.

Livermore, D. M. (1987). Clinical significance of beta-lactamase induction and stable derepression in gram-negative rods. Eur. J. Clin. Microbiol. 6, 439–445. doi:10.1007/BF02013107.

Michniewski, S., Redgwell, T., Grigonyte, A., Rihtman, B., Aguilo-Ferretjans, M., Christie-Oleza, J., et al. (2019). Riding the wave of genomics to investigate aquatic coliphage diversity and activity. Environ. Microbiol. 21, 2112–2128. doi:10.1111/1462-2920.14590.

Montso, P. K., Mlambo, V., and Ateba, C. N. (2019). Characterization of Lytic Bacteriophages Infecting Multidrug-Resistant Shiga Toxigenic Atypical Escherichia coli O177 Strains Isolated From Cattle Feces. Front. Public Heal. 7. doi:10.3389/fpubh.2019.00355.

Nale, J. Y., Spencer, J., Hargreaves, K. R., Buckley, A. M., Trzepinski, P., Douce, G. R., et al. (2015). Bacteriophage Combinations Significantly Reduce Clostridium difficile Growth In Vitro and Proliferation In Vivo. Antimicrob. Agents Chemother. 60, 968–981. doi:10.1128/AAC.01774-15 [doi].

Peirano, G., Chen, L., Kreiswirth, B. N., and Pitout, J. D. D. (2020). Emerging Antimicrobial Resistant High-Risk clones among Klebsiella pneumoniae1: ST307 and ST147. Antimicrob. Agents Chemother. doi:10.1128/aac.01148-20.

Rivera, D., Hudson, L. K., Denes, T. G., Hamilton-West, C., Pezoa, D., and Moreno-Switt, A. I. (2019). Two phages of the genera felixunavirus subjected to 12 hour challenge on salmonella infantis showed distinct genotypic and phenotypic changes. Viruses 11. doi:10.3390/v11070586.

Schmerer, M., Molineux, I. J., and Bull, J. J. (2014). Synergy as a rationale for phage therapy using phage cocktails. PeerJ 2014. doi:10.7717/peerj.590.

Seemann, T. mlst, Github. Available at: https://github.com/tseemann/mlst [Accessed February 19, 2020].

Seemann, T. (2014). Prokka: Rapid prokaryotic genome annotation. Bioinformatics 30, 2068–2069. doi:10.1093/bioinformatics/btu153.

Storms, Z. J., Teel, M. R., Mercurio, K., and Sauvageau, D. (2019). The Virulence Index: A Metric for Quantitative Analysis of Phage Virulence. PHAGE 1, 17–26. doi:10.1089/phage.2019.0001.

Sybesma, W., Zbinden, R., Chanishvili, N., Kutateladze, M., Chkhotua, A., Ujmajuridze, A., et al. (2016). Bacteriophages as potential treatment for urinary tract infections. Front. Microbiol. 7. doi:10.3389/fmicb.2016.00465.

Sykes, R. B., and Matthew, M. (1976). The beta-lactamases of gram-negative bacteria and their role in resistance to beta-lactam antibiotics. J. Antimicrob. Chemother. 2, 115–57. doi:10.1093/jac/2.2.115.

Tabassum, R., Shafique, M., Khawaja, K. A., Alvi, I. A., Rehman, Y., Sheik, C. S., et al. (2018). Complete genome analysis of a Siphoviridae phage TSK1 showing biofilm removal potential against Klebsiella pneumoniae. Sci. Rep. 8, 17904–17911. doi:10.1038/s41598-018-36229-y.

Taha, O. A., Connerton, P. L., Connerton, I. F., and El-Shibiny, A. (2018). Bacteriophage ZCKP1: A Potential Treatment for Klebsiella pneumoniae Isolated From Diabetic Foot Patients. Front. Microbiol. 9, 2127. doi:10.3389/fmicb.2018.02127.

van Duin, D., and Doi, Y. (2017). The global epidemiology of carbapenemase-producing Enterobacteriaceae. Virulence 8, 460–469. doi:10.1080/21505594.2016.1222343 [doi].

Zankari, E., Hasman, H., Cosentino, S., Vestergaard, M., Rasmussen, S., Lund, O., et al. (2012). Identification of acquired antimicrobial resistance genes. J. Antimicrob. Chemother. 67, 2640–2644. doi:10.1093/jac/dks261.

