## Supplementary Data S7 for "Analysis of selection methods to develop novel phage therapy cocktails against antimicrobial resistant clinical isolates of bacteria"

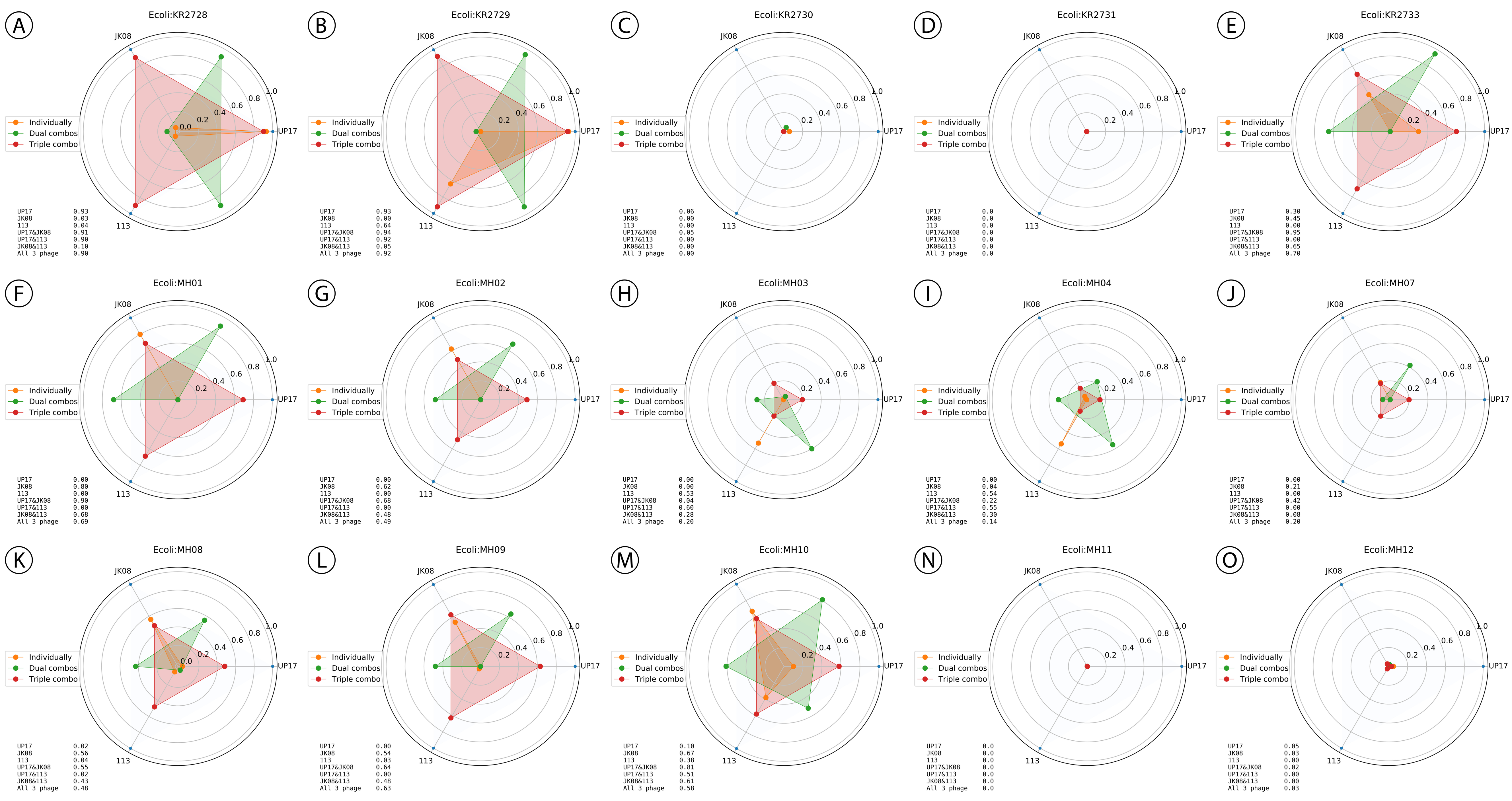

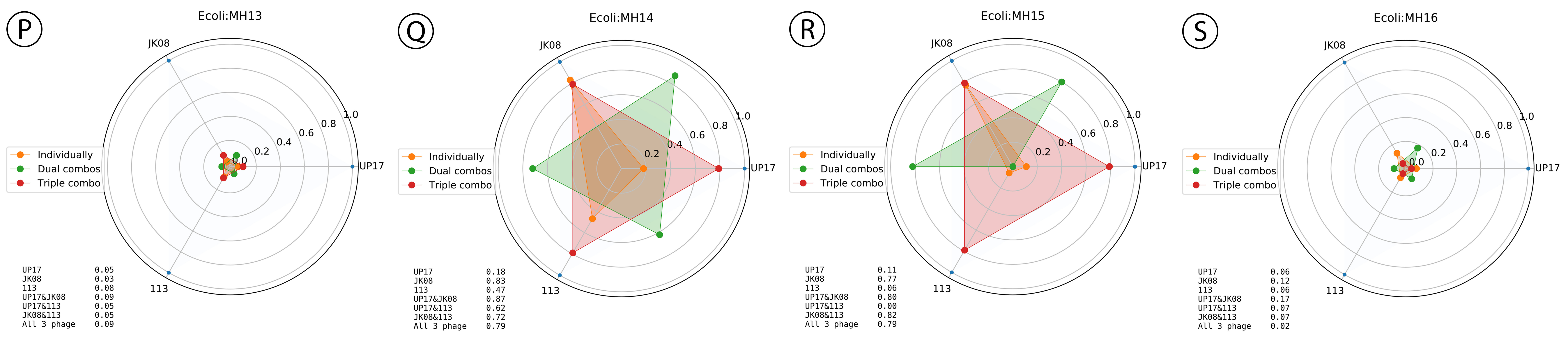


Figure S7 A spider diagram depicting the virulence index scores of individual phage and various phage combinations across all 19 ESBL-producing E. coli. Each clinical isolate is represented by a circle, this circle is divided into smaller circles with the outer circle representing a virulence index score of 1 and the centre of the large circle representing 0; each smaller circle representing 0.2 on the scale. Each of the three phages (UP17, JK08 and 113) are labelled on the outer circle with a representative line to the centre of the circle. The coloured dots represent different combinations of phages (orange – individual phage, green – doublet combination, red - triplet combination).
